# Morphodynamics of tip growing cells

**DOI:** 10.1101/2024.11.17.624014

**Authors:** Gareth Wyn Jones, Otger Campás, L. Mahadevan

## Abstract

Pollen tubes, root hairs and fungal hyphae elongate by adding material at their tips. The region in the immediate vicinity of the tip is dynamically pliable and allows for the addition of new wall material even as it flows in response to the turgor that drives it. Any imbalance between the rate of material addition and its consequent flow will cause a tip growing cell to either burst because its wall thins or stop growing because its wall thickens. We use an experimentally-inspired feedback law that couples vesicle exocytosis to wall mechanics via the local strain rate to construct a minimal theory for the dynamics of tip growing cells. Our theory characterizes the parameter regime where stable steady tip growth is possible in terms of two dimensionless parameters: a scaled turgor pressure, and a ratio of length scales describing gradients of wall extensibility and vesicle composition near the tip. Our analysis also explains the experimentally observed shape response of tip growing cells observed when external turgor is dynamically varied. All together our formalism provides a general framework for the coupled dynamics of internal vesicular transport and wall mechanics in tip-growing cells.

## INTRODUCTION

Unicellular organisms have a wide variety of shapes and sizes and can grow and change shape dynamically. They use this ability to migrate, expand into their environment, or change the shape of their host tissue. Cell shape is controlled by physical mechanisms such as material properties, mechanical forces, and mass transport. These factors are linked together in a complex network of interactions, under strong genetic control — though the genes do not dictate the cell morphology directly. This network of interactions is poorly understood in general, and in particular it is not clear how the range of morphologies available to the cell is constrained by the physics of the cell shaping process, although recent work in prokaryotic cells such as bacteria has begun to unravel the physical basis for cell shape by focusing on the role of wall mechanics, maintenance and internal turgor [1]. Here we build on previous work that focused on the steady growth of cells [2] and its comparative dynamics [3] and demonstrate how the morphology of a certain class of elongating cells can be determined and controlled dynamically by its physical configuration: namely the spatial distribution of its mechanical properties and its secretion profile.

Many studies aimed at understanding the factors governing cell shape have focused on intracellular signaling pathways that interact with the cytoskeletal network [4, 5], or document the shape changes arising from specific gene mutations [6, 7]. The role of mechanics in cell morphology is well-known [8], and research in many different systems has attested to its importance [9–11]. Organisms whose cells possess a stiff cell wall, including plants, fungi, and certain bacteria, have proven particularly advantageous for studying cell growth and morphogenesis, since the cell shape is easily measured by the extent of the wall. The main driver of shape change in such cells is mechanical, since high stresses are required to overcome the stiffness of the cell wall, and deformations tend to progress at low strain rates, again aiding the shape measurement. One model system for growth in walled cells is the class of tip-growing cells. These grow as long filaments with a dome-shaped tip; the cells’ shape is maintained by confining their growth to the tip region, thus elongating the cell in a self-similar fashion [12]. Such tip-growing cells include fungal hyphae [13], water molds [14], rhizoids and protonemata in primitive plants such as mosses and ferns [15], and the pollen tubes and root hairs of seed plants [16–18].

All walled cells possess a strong internal turgor pressure, generated by the osmotic flow of water into the cell interior. While this pressure is the ultimate driver of cell growth, it is an isotropic quantity and thus cannot control the directionality of growth — nor the tip morphology — by itself [17, 19]. This directionality is provided by a selective weakening of the cell walls at the elongating tip. As the material of the wall is weakened, it can no longer withstand the turgor pressure, and is forced to expand. In tip-growing cells this process occurs at the apex, which drives through the extracellular medium leaving an elongated cylindrical filament in its wake [12]. Without a constant flow of new cell wall material to the growing region, the wall would become increasingly thinner, and eventually burst. The new material is either transported to or assembled at the growing tip. The specific molecular machinery and cellular process that accomplish this are different across species that span pollen tubes, yeast, fungi and molds. This material comes packaged in vesicles, which also include enzymes that soften the wall material [13, 20–22]. In some cases the vesicles’ contents are incorporated into the cell wall by exocytosis, with their membranes providing the necessary additional material for the expansion of the cell’s lipid bilayer membrane. In other cells, such as yeast, exocytosis is used to deliver cell wall synthases that then assemble the cell wall directly from the cell surface [23].

Modeling studies have become a key tool in determining how the characteristic behavior of tip-growing cells emerges from their mechanical and biochemical properties [24, 25]. Many models of tip growth have focused on the dynamics of vesicle distribution [26], how this distribution influences cell shape [27, 28], or on the process of vesicle exocytosis [29]. These studies confirm the importance of vesicle dynamics in tip growth, but fail to couple this to the mechanics of the cell wall. Conversely, mechanical models of the cell wall, [2, 30–32], tend to neglect the exocytosis process by which new material is added to the wall. More recently, some models have coupled the cell wall mechanics to cell wall assembly via mechanical feedbacks in budding [23, 33] and fission yeast [34]. However, the mechanical feedbacks are genetically encoded and very specific to the organisms. It is unclear if more generic couplings between cell wall mechanics and assembly via exocytosis exist across species. Recent theoretical studies coupled the dynamics of the plasma membrane and the expanding cell wall, but only in a 1D flat geometry [35]. More recently, it has been experimentally shown that membrane dynamics is essential for cellular morphogenesis [36]. Therefore, coupling the membrane dynamics to mechanics of the expanding cell wall in the cell’s curved 3D geometry is necessary to account for the cell morphogenesis. Modeling the dynamics of plasma membrane transfer between the vesicles and cell tip introduces to the model coupling between cell wall mechanics and the secretion of new cell wall material, which is critical to understanding the dynamics of the overall system.

Observations of tip-growing cells have uncovered a number of testable phenomena, including the tendency of such cells to cease growth as the turgor pressure drops below a certain value — though the thickness continues to increase [37]. Notably this effect is reversed if the turgor pressure is restored [38]. As a further example, pollen tubes have been shown to have an elevated region of exocytosis in an annular region surrounding the tip of the cell [20]. These observations, together with the wide variety of observed cell shapes, should be accessible to a computational model by variation of input parameters.

The model presented here explicitly accounts for the conservation of plasma membrane through the exocytosis and endocytosis of vesicles, which is coupled to a fluid mechanical model of the cell wall through a term embodying the addition of new wall material. The effect of turgor pressure, vesicle distribution and wall rheology can be embodied in three dimensionless parameters, which are sufficient to explain many experimental observations of tip-growing cell behavior, including those indicated above.

## METHODS

Our primary assumption is that the tip-growing cell has two components: a lipid bilayer membrane, enclosed by a viscous fluid sheet representing the cell wall. The viscous sheet flows under the influence of internal pressure, and transport vesicles bring new wall material and membrane to the growing cell. Being thin, the geometry of the fluid sheet can be reduced to that of its middle surface, whose natural coordinates are the arc length *s* measured from the tip, and the azimuthal angle *ϕ* (see the Supplementary Material for details of the modeling). As the shape will be assumed to be axisymmetric, changes in the azimuthal direction will be neglected. The shape of the middle surface is described by two variables *r* and *θ*, both functions of *s* (and time, *t*), linked through *∂r/∂s* = cos *θ*, as shown in Fig. 1(a). Using these one may derive the principal curvatures κ_1_ = *∂θ/∂s*, κ_2_ = *r*^−1^ sin *θ* of the middle surface.

**FIG. 1.**
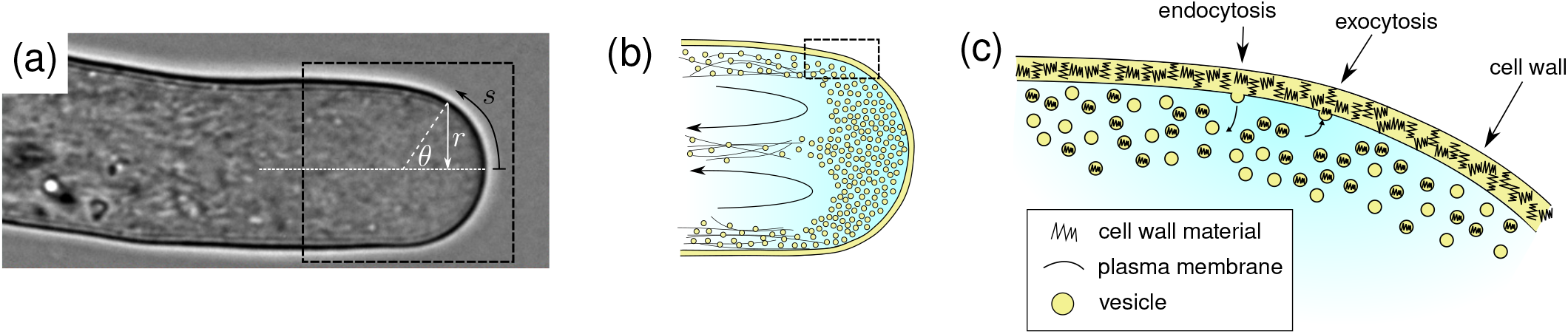
Morphology of a tip-growing cell and coordinate representation. The angiosperm pollen tube, an archetypal tip-growing cell. (a) An image of a lily pollen tube, reproduced from Campás et al. [2], overlaid with the geometric parameters *s, r* and *θ*. (b) Vesicle dynamics and distribution. Vesicles are transported towards the tip by an actin–myosin cytoskeletal network and collect in a reservoir at the tip. Excess vesicles are transported back towards the shank of the tube along the center line. (c) Close-up of the exo- and endocytosis process at the cell wall: exocytosis adds cell wall material and plasma membrane; endocytosis removes plasma membrane.

We now consider the equation governing the change in plasma membrane area under exocytosis. Suppose that the number of vesicles exocytosed per unit area per unit time (the exocytosis rate) at a location *s* is denoted *Q*(*s*). We similarly define an endocytosis rate *R*(*s*). Furthermore, suppose that the exocytosing vesicles carry on average a mass *m*_*w*_(*s*) of cell wall precursor material and a mass *m*_*m*_(*s*) of membrane material. Conversely, assume that no wall material is contained within endocytosing vesicles and that they only contain (again, on average) a mass *m*_*m*_(*s*) of membrane material. Consider a small patch of cell wall of area Δ*A*. The net increase in membrane mass here through exo- and endocytosis in a time Δ*t* is

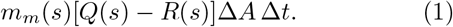

Due to the motion of the fluid sheet, the cell wall area Δ*A* increases by an amount 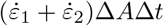 in the same period, where 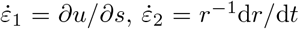 are the strain rate components in the cell wall; *u*(*s, t*) is the cell wall velocity. Thus the additional membrane mass required to keep the amount of membrane per unit cell wall area constant is

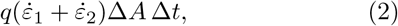

where *q* is the area density of membrane material. Under the assumption that this homeostatic property is attained by the cell, we find

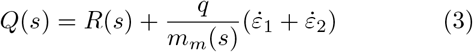

by equating Eq. 1 and Eq. 2 which couples the mechanics of the cell wall and exo- and endo-cytosis in a generic manner.

The rate of increase in cell wall material mass per unit area is *γ* = *m*_*w*_*Q*, or

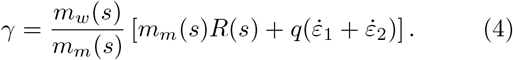

This term appears in the equation for the conservation of cell wall mass,

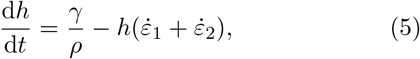

where *h*(*s, t*) is the thickness of the cell wall and *ρ* is the density of cell wall material. Through these two equations, the vesicle secretion dynamics and the cell wall dynamics are coupled, and both processes need to be accounted for in order to understand the cell growth.

The system is closed by equations describing the mechanics of the fluid sheet. These can be derived from physical principles [2] or systematically from the equations of fluid mechanics [39]. We choose a fluid-based model for the cell wall mechanics because it is able to capture the conservation of mass following vesicle exocytosis [2]; elastic-based models typically require the selection of constitutive assumptions to achieve this consistently. The mechanical equations embody a balance of forces in the cell wall, and a constitutive law asserting the linear dependence of stress on strain rate (Supplementary Material). The key quantity in the mechanical description is the cell wall viscosity *µ*(*s*), which characterizes the extensibility of the wall; we take this to be an input of the system.

Combining all previous equations and performing a nondimensionalization that scales all variables with physically-relevant quantities (see the Supplementary Material), we obtain the following system of equations (for dimensionless quantities 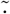):

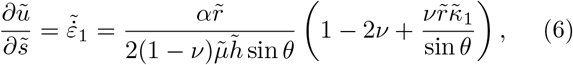

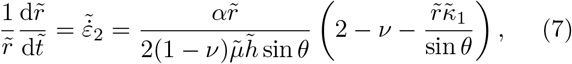

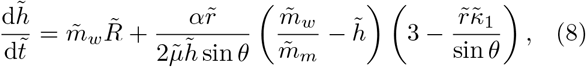

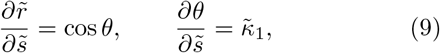

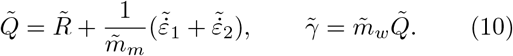

In this formulation, *ν* refers to the Poisson ratio analog of the fluid (taken to be 1*/*2 for an incompressible fluid), and the dimensionless parameter *α* is given by

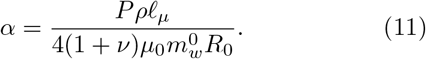

This formula incorporates the turgor pressure *P* in the cell, typical scalings for viscosity *µ*_0_ and vesicle dynamics 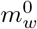 and *R*_0_, and a typical length scale 𝓁_*µ*_ of the problem (which we choose to be that of spatial variations in viscosity). High values of *α* correspond to high turgor pressure, a low viscosity, or a low mass addition rate, and can therefore be expected to result in a thinner and more rapidly-expanding cell wall, with the opposite behavior as *α* → 0.

Though *α* is the only dimensionless parameter appearing explicitly in the equations, there may be further dimensionless parameters hidden in the expressions for 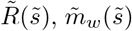 and 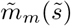, which may have different length scales. To investigate how these relative length scales affect the model, we select profiles for these quantities as model inputs. We choose the following dimensional forms for the input quantities:

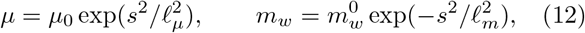

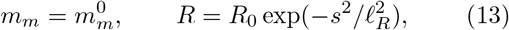

where 𝓁_*µ*_, 𝓁_*R*_ and 𝓁_*m*_ are length scales associated with gradients in the viscosity, endocytosis rate, and cell wall material per vesicle respectively near the tip (note that we normalize all length scales with that of viscosity).

These functional forms (illustrated in Fig. 2) encapsulate observed features of tip-growing cells in the *s* → ∞ limit, from the tip down the shank. The viscosity *µ* increasing to infinity models the hardening of the cell wall in this limit due to increased cross-linking of cell wall components [2]; cell wall material per vesicle *m*_*w*_ (henceforth “vesicle composition”) decreasing to zero models the fact that vesicles containing cell wall material are generally delivered to the tip [15]; and a decreasing endocytosis rate *R* reflects the observation that endocytosis is primarily observed in the tip [20]. The remaining parameter *m*_*m*_ is set to a constant for simplicity.

**FIG. 2.**
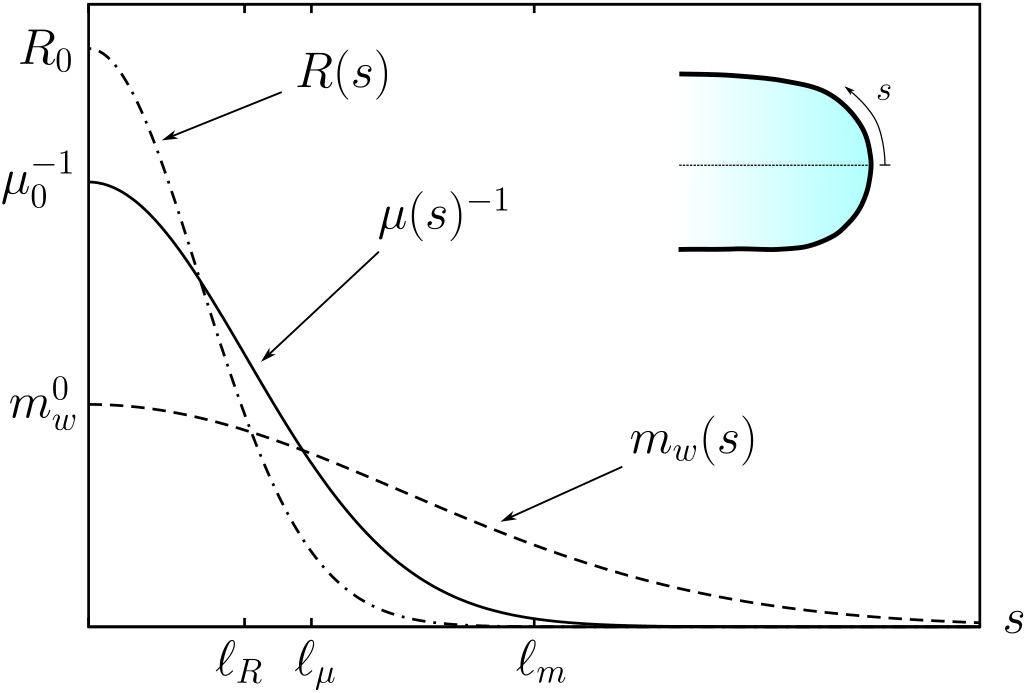
Distribution of parameters with arc length. A representative diagram demonstrating how varying the three length scales 𝓁_*µ*_, 𝓁_*m*_, and 𝓁_*R*_ affect the three spatially-varying quantities of fluidity (inverse viscosity) *µ*(*s*)^−1^, quantity of cell wall material within vesicles *m*_*w*_ (*s*), and endocytosis rate *R*(*s*) respectively. The forms of each quantity are given in Eq. 12–13.

Rewriting Eq. 12–13 in dimensionless form yields

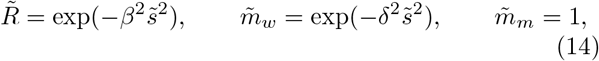

leading to two additional dimensionless constants:

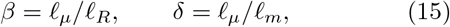

that correspond to the ratios of the length scales of viscosity and endocytosis, and viscosity and vesicle composition, respectively. If *δ* ≫ 1, the viscosity length scale is larger than that of the vesicle composition, leading to a region of the cell wall where little new mass is being added but the material is still quite fluid. Conversely, if *δ*≪ 1 then the composition of the vesicles remains largely constant over the region where the viscosity increases to infinity.

The system in Eq. 6–10 when discretized in space yields a differential–algebraic equation system [40], which we solve using the method of lines. For full details, refer to the Supplementary Material.

## RESULTS AND DISCUSSION

The behavior of solutions to the system in Eq. 6–10 is governed by the parameters *α, β* and *δ*. Both *β* and *δ* primarily affect the spatial decay of the mass addition rate *γ*, and so we decoupled these parameters, first by varying *δ* and letting *β* = 0 (so that 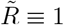), and then by varying *β* and keeping 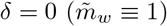.

In the first calculation we sought the steady solution to Eq. 6–10 for a set of parameters *α, δ* in the range 0.1 to 10. The nature of the solutions for each parameter choice is plotted logarithmically in Fig. 3(a). On the left-hand side of the figure is a region where no physical steady solution was found: the wall continued thickening as the time increased. At the same time, the velocity tended to zero, so this region corresponds to the case where the cell elongation ceases.

**FIG. 3.**
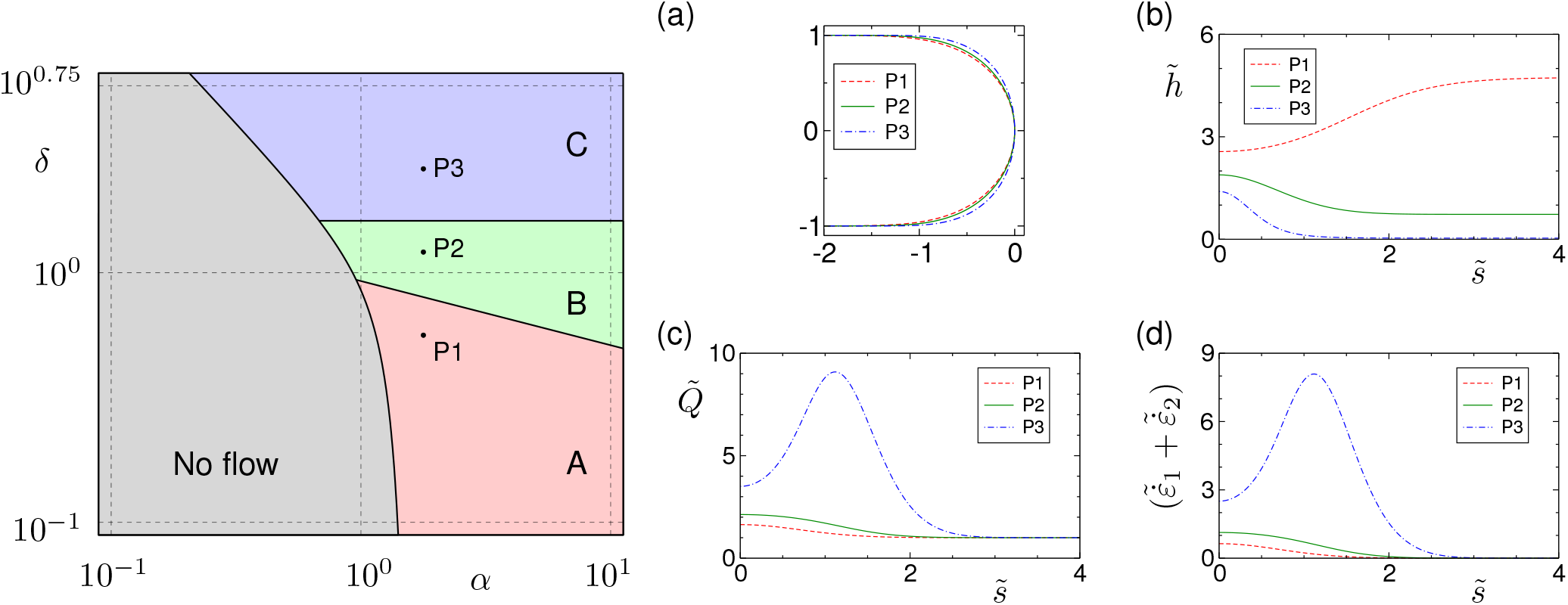
Phase diagram of morphologies as a function of the dimensionless turgor pressure *α* and the ratio of spatial distributions *δ* defined in Eq. 15. Parameter sweep of *α* against *δ* based on 550 individual calculations, displaying behavior of the stationary solution to the tip growth equations in Eq. 6–10 when *β* = 0. Stationary solutions are displayed for three parameter values: *α* = 1.78 and *δ* = 0.56, 1.21, 2.61, which are shown on the *α*–*δ* map. (a) The shape of the cell tip, scaled so that 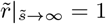. (b) Cell thickness. (c) Exocytosis rate. (d) Strain rate of the cell wall.

A steady solution was found in the rest of the parameter space, which can be subdivided into three regions A, B and C, depending on the nature of the strain rate and thickness (see Table I). These regions correspond to different orders of magnitude of the parameter *δ*. In Fig. 3(b)–(d) we plot some typical solutions, for *α* = 1.78 and for five values of *δ*. We can see the behavior of strain rate and thickness as described in Table I, together with the differences in cell shape and cell wall growth rate 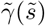.

**TABLE 1.** Characterization of the three regions A, B, C in Fig. 3 based on the strain rate, thickness, and magnitude of *δ*.

| Region | Max strain rate at tip | $\tilde{h}_{\text{tip}} > \tilde{h}_{\text{shaft}}$ | $\delta$ |
| --- | --- | --- | --- |
| A | Yes | No | $\delta \ll 1$ |
| B | Yes | Yes | $\delta \sim 1$ |
| C | No | Yes | $\delta \gg 1$ |

Comparing solutions as *α* varies and *δ* remains constant, there is little change in the shape of the cell (see Supplementary Material). The only significant trend is that as *α* increases, the strain rate (and velocity and growth rate) increases and the thickness decreases.

On allowing 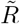 to vary while keeping 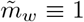, we find that the *α*–*β* parameter space is similar in structure to the *α*–*δ* parameter space above (details can be found in the Supplementary Material).

Eq. 8 indicates that two terms contribute to the wall growth rate: a term linearly dependent on the strain rate (and independent of 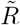), and a strain-independent term 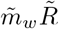. Since allowing spatial variations in both 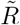 and 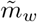 leads to the same basic structure of parameter space, we conclude that the strain-independent, or ‘baseline’ mass addition term 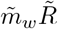 determines the structure, and we can interpret the three regions A, B and C in terms of the different patterns of mass addition prescribed by this term.

In region B, the region of mass addition and the region of fluidized cell wall have approximately the same extent; this is the most common regime observed experimentally. Both the thickness 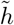 and the strain rate have their maxima at 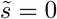, at the cell tip.

In region A, the region of mass addition is longer than the region of low viscosity, resulting in a part of the cell where the wall has largely solidified but new material is still being added. The result of this mass addition is that the wall thickness increases, and the strain rate is considerably smaller than in region B.

In region C, we see the opposite effect. There is a region of the cell wall which has relatively low viscosity, but with a much smaller rate of mass increase. This means that the higher fluid velocity due to the low viscosity is not balanced by a thickening from mass addition, and so the strain rate is increased in this location and the wall thickness is smaller. Furthermore, there is a peak in the curvature 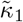 near the peak in strain rate, indicating that the tip geometry is flattened, in contrast to the pointed morphology in regions A and B. Note that the maximum strain rate becomes ever larger as *β* and *δ* increase (significantly so in the case of large *δ*), to a point at which the model breaks down and the cell can be seen as having burst. This is consistent with a cell wall piercing regime previously reported [23]. In all cases the extent of the flowing region of the cell tip is determined by the length scale of viscosity, which in these dimensionless variables is kept at unity.

The existence of the no-solution region can be understood by considering the limit as *α* → 0, when the equation for the thickness reduces to 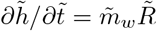, and thus the thickness increases without bound. To counteract this there must be a minimum value of *α* which provides just enough flow to overcome this thickening tendency. For high values of *δ* or *β*, the flow generated by a given value of *α* is higher, and so the minimum value of *α* necessary to produce a steady flow is lower.

Though the results show that the mass addition rate *γ* into the cell wall is always at its maximum at the tip, the two experiments (*β* = 0, *δ* = 0) lead to different forms of the exocytosis rate *Q*. In the former case, 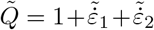, and so, the peak in *Q* is coincident with the peak in strain rates (which in area C is in an annular region away from the tip). In the latter case, 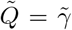 and hence the exocytosis rate is always monotonically decreasing from the tip. We expect the result of both *β* and *δ* being nonzero to be intermediate between these two extremes.

### Cell shape dependence on functional forms of material addition and viscosity

The choice of decay profile for the quantities 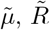 and 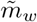 up to this point has been chosen to be of an exponential form. This leads to modest shape changes under certain parameter variations. A more dramatic shape change can be shown if we allow these quantities to decay as a power law as 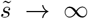, e.g. 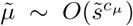 and 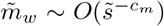 (see Supplementary Material for details; a choice of *c*_*m*_ = corresponds to exponential decay). Then, on choosing some representative values *α* = 2, *β* = 0, *δ* = 1, we can plot solutions for different combinations of *c*_*µ*_, *c*_*m*_.

Though the shape appears to remain invariant under variations in the decay of 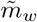, a more gradual solidification of the cell wall clearly leads to more pointed morphologies, as shown in Fig. 4 over different values of *c*_*µ*_. We conclude that the nature of the solidification as 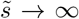 is a key parameter in determining the shape of the tip.

**FIG. 4.**
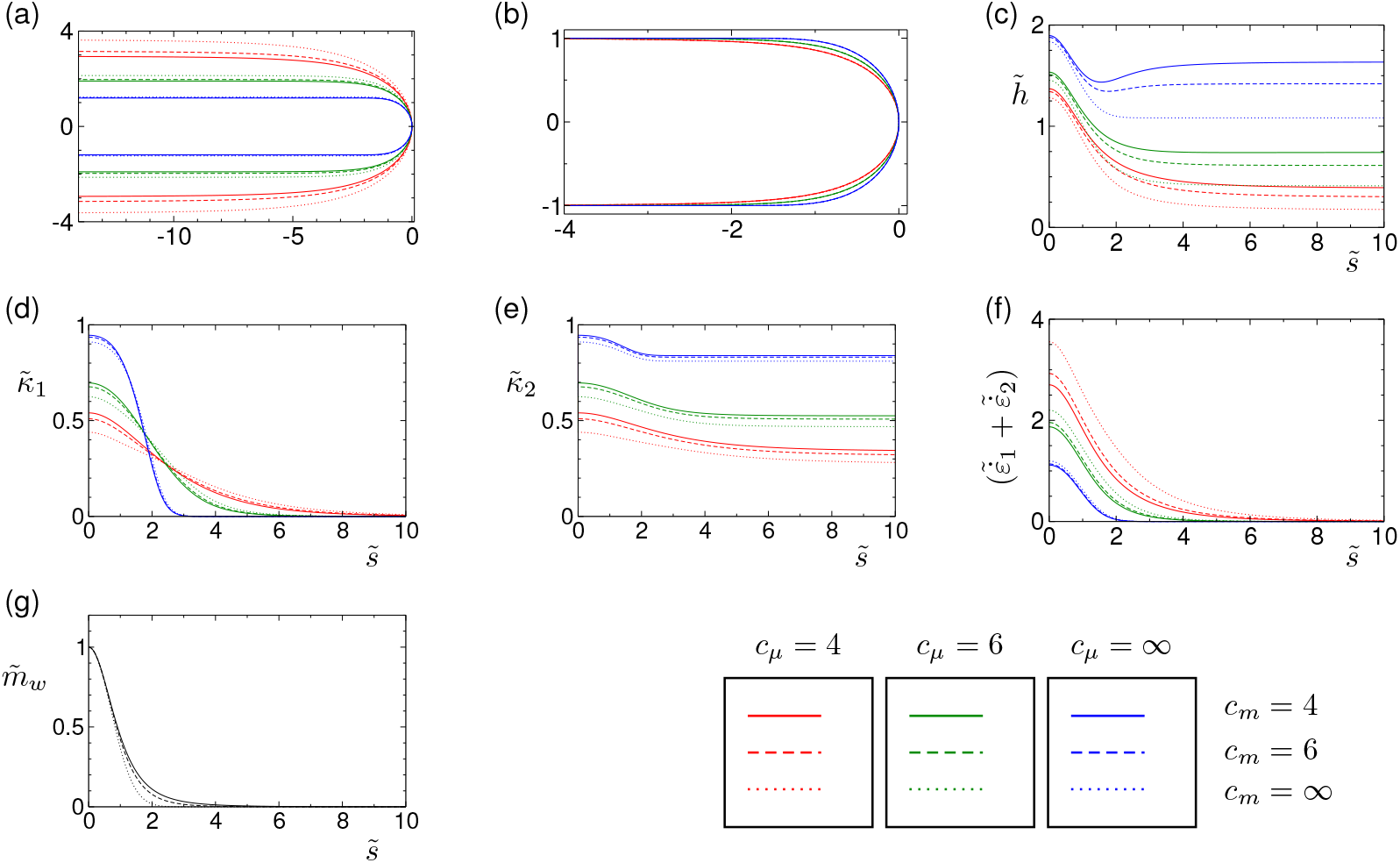
Role of functional forms of wall addition rates and viscosities. Stationary solutions for *α* = 2, *β* = 0, *δ* = 1, for power-law forms of the wall addition rates and viscosities (see Supplementary Material for details) and the nine combinations of *c*_*m*_ ∈ {4, 6, ∞} and *c*_*µ*_ ∈ {4, 6, ∞}. (a) Tip shape in the original dimensionless variables. (b) Tip shape, scaled so that the shank is of radius 1. (c) Cell wall thickness. (d)–(e) Principal curvatures of the cell wall. (f) Wall strain rate. (g) The distributions of 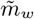; the distributions of 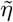 are coincident. Bottom right shows the color codes for the choices of the power law forms.

### Time-dependent solutions

The time-dependent description can be tested in an experimental situation that involves temporal changes in the dynamics of tip growth. In the experiments by Zerzour et al. [38], the tube elongation rate was arrested through a reduction of turgor pressure, with the thickness continuing to grow in the tip. As the turgor pressure was restored to its original value, the growth recommenced, with the additional material causing a thick bulge in the shank as it continued to elongate. We have reproduced this experiment in Fig. 5, and show that this behavior can be interpreted in terms of the *α*–*δ* parameter space (and a similar effect would be seen were we to consider a spatially-varying 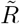). The lowering of turgor pressure corresponds to a lowering in *α*, causing the system to leave the stable region at the right and enter the “no flow” region to the left (see Fig. 5(b)). In this region, as noted earlier, strain rate falls to zero and the thickness continues to rise as a result of the baseline mass addition term 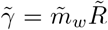. Tube elongation is recovered by restoring turgor pressure (or *α*), and the bulge is retained by the fluid sheet solidifying as it lengthens.

**FIG. 5.**
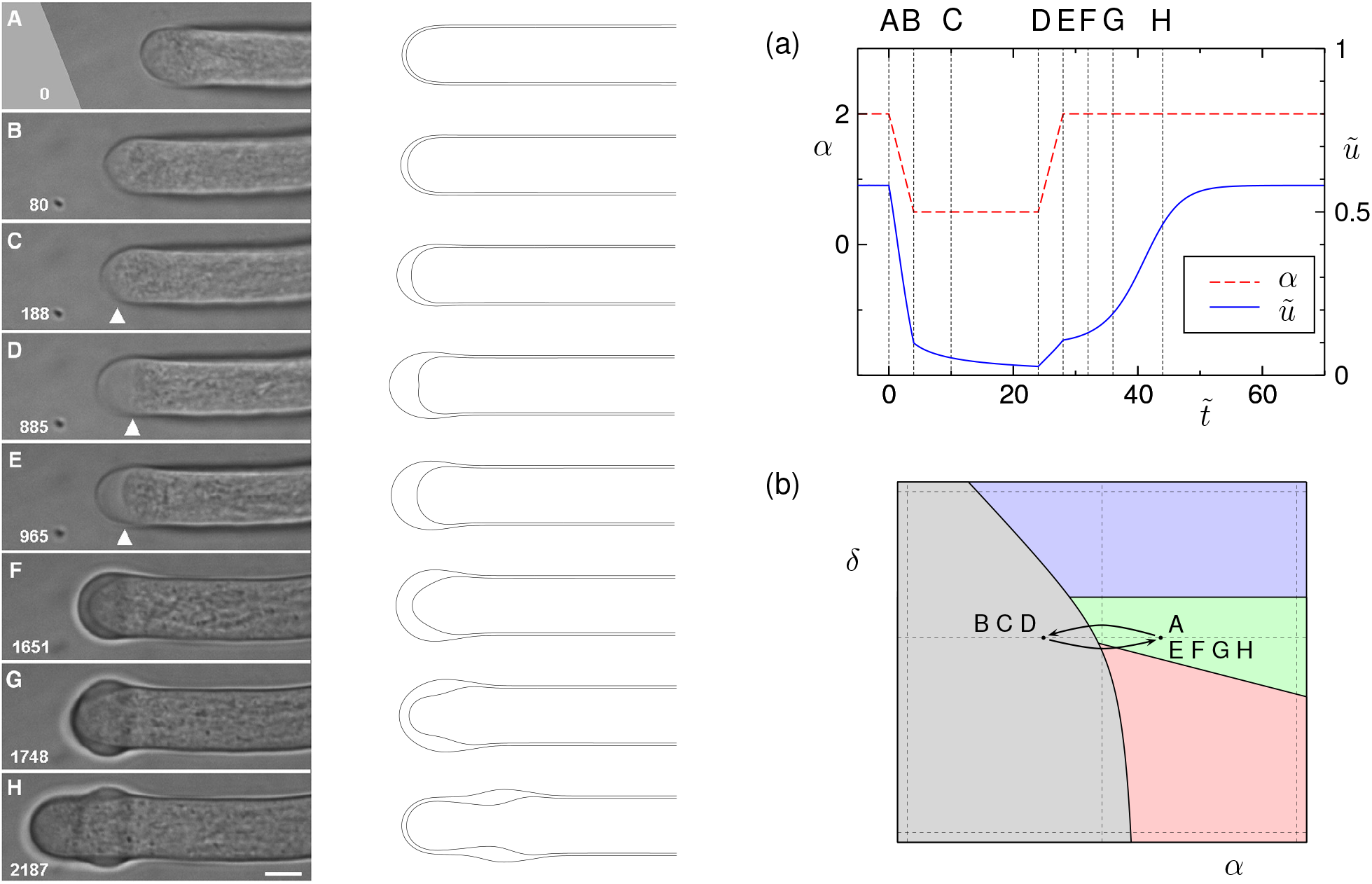
Dynamical transitions of wall morphologies when external turgor is changed. Left: images of an experiment where a growing pollen tube has its turgor pressure lowered, causing growth to cease and a gradual thickening of the tip; pressure recovery causes growth to recommence. Reproduced from Zerzour et al. [38]. Center: a numerical recreation of the experiment with *β* = 0 and *δ* = 1; lowering turgor pressure causes the elongation rate to cease and the tip to thicken. Turgor pressure recovery causes growth to recommence and the thickened region to be advected down the tube shank. (a) Profile of the turgor pressure *α* and elongation rate 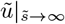. (b) Interpretation of the experiment: the turgor pressure is lowered from a steady-solution region into the no flow region, and back again.

## CONCLUSION

Our theory shows how the shape and dynamics of tip-growing cells emerges from the interplay of mechanics and secretion by coupling mass and force balance terms in the cell wall to the conservation of plasma membrane through vesicle secretion. Our analysis of the possible dynamical scenarios have shown that a wide range of behaviors and morphologies can be predicted in terms of the physical and mechanical inputs to the model.

Notably, we predict that a lowering of turgor pressure is sufficient to cause cell growth to cease, though the wall continues to thicken. This is a well-established experimental observation [37, 38], but mechanical models to date have either failed to capture the behavior or have interpreted it as evidence for a yield stress in the cell wall material. Our results show that resorting to such complex rheological models is unnecessary, and that the phenomenon can simply arise from a combination of persistent material addition and a turgor pressure which is too small to compensate. In our model this behavior is captured by lowering the value of *α*; it would be instructive to observe whether experimentally increasing cell wall viscosity or endocytosis rate (which, through Eq. 11, also reduce *α*) have a similar growth-cessation effect. Increasing *α* in our model leads to a thinner and more rapidly deforming cell wall. This agrees with observations that the cell wall ruptures at the tip if the turgor pressure *P* is elevated past a critical value.

A second result of our model is that for values of *β* or *δ* which are much greater than 1 (where the viscosity length scale is much longer than the region of mass addition), the strain rate reaches a peak in an annular region some distance from the tip of the tube. The existence of this annular region has been observed in root hairs [41]. In the case of *β* = 0 (constant endocytosis) and large *δ*, this region of high strain rates leads to a region of high exocytosis *Q*, due to the requirement that the area of plasma membrane is conserved. The presence of such an annular peak in exocytosis has also been inferred from recent experimental observations [20]. Our results do not show a corresponding peak in the rate of cell wall growth *γ*, though this may change if non-monotonic profiles of *m*_*w*_, *R* or *µ* were selected.

Thirdly, we have shown that variations in model parameters lead to changes in the shape of the tip. In particular, *c*_*µ*_, the nature of the increase in viscosity as the wall hardens, has a noticeable effect on morphology. As the wall hardens with a more gradual power-law behavior, the cell tip adopts a sharper profile. We envisage that this result could lead to new insights into the nature of the hardening behavior of cell walls based on observed variations in profile shapes between different species [3, 42, 43].

In our model, *m*_*m*_, *m*_*w*_, *R* and *µ* have been prescribed independently as input variables. In reality, the profiles of these quantities will emerge as a result of complex interactions with signaling networks, cell biochemistry, and cytoskeletal dynamics. Ultimately, genetic variation causes changes in cell morphology by modulating this meshwork of interactions to produce different forms of the input variables, but the universe of shapes is eventually constrained by mass balance and force balance. The model presented here is simple and flexible enough that the interactions between the input variables can be incorporated through the introduction of additional equations. Even in the absence of such dynamic relations, our model can verify purported mechanisms of tip growth morphogenesis based on experimentally-measured profiles of these four parameters.

## AUTHOR CONTRIBUTIONS

L.M. and O.C. conceived of the study. G.J., O.C. and L.M. developed the theoretical model, G.J. performed the simulations of the model and G.J., O.C. and L.M. analyzed the results. G.J., O.C. and L.M. wrote and edited the manuscript.

## ACKNOWLEDGMENTS

This work was partially supported by the Deutsche Forschungsgemeinschaft (DFG, German Research Foundation) under Germany’s Excellence Strategy – EXC 2068 – 390729961– Cluster of Excellence Physics of Life of TU Dresden (OC), the Simons Foundation(LM), the MacArthur Foundation(LM) and the Henri Seydoux Fund (LM).

